# Efficient estimation of time-dependent functional connectivity using Structural Connectivity constraints

**DOI:** 10.1101/2022.09.21.508847

**Authors:** Hernan Hernandez Larzabal, David Araya, Lazara Liset Gonzalez Rodriguez, Claudio Roman, Nelson Trujillo-Barreto, Pamela Guevara, Wael El-Deredy

## Abstract

Multivariate autoregressive models [MAR] allows estimating effective brain connectivity by considering both power and phase fluctuations of the signals involved. A MAR models brain activity in one region as a linear combination of past activations in all other regions. A Hidden Markov model, HMM, whose states’ emisions are drawn from state-specific MARs, can then be used to model fast switching of effective brain connectivity over time. However, the large number of MAR parameters, impede the accurate and efficient estimation of such models from neuroimaging timeseries with limited length.

We propose a new model for inferring time-dependent effective brain connectivity by using a sparse MAR parameterisation to model the states’ emisions of a Hidden Semi-Markov Model, HsMM-MAR-AC. The sparse MAR model parameters are restricted by Anatomical Connectivity information in two ways: direct effective connectivity between two regions is only considered if the corresponding structural link/connection exists; and the time-lag associated with each direct connection is computed based on the average fibre length between the two regions, such that only one lag per connection is estimated.

We simulated ground truth time-dependent brain connectivity states by generating time-series of 4 to 10 minutes sampled at 5ms, from switching Resting State Networks with a reference structural connectivity, and evaluated the accuracy of the new model in recovering the simulated Brain connectivity States against different levels of connectivity thresholding above and below the reference.

We show that even when restricting the MAR to half of the reference connections, the model was able to recover the number of brain states, the associated connectivity features and their dynamics, with as little data as 4 mins. More relaxed structural connectivity thresholds required longer data to estimate the model accurately, and became computationally unfeasible without anatomical restrictions.

HsMM-MAR-AC offers an efficient algorithm for estimating time-depend Effective Connectivity (tdEC) from neuroimaging data, that exploits the advantages of MAR without identifiability problems, excessive demand on data collection, or unnecessary computational complexity.

## 1 INTRODUCTION

A current challenge in the study of the brain is the detection of brain states (BS) transitions, either in resting condition or evoked by task using EEG/MEG (Olier et al. (2013), Trujillo-Barreto et al. (2019), Vidaurre et al. (2018),Khanna et al. (2015),O’Neill et al. (2018),Woolrich et al. (2013)). This challenge is known as the BS allocation problem. Here a BS corresponds to an operational mode of the brain that produces a quasi-stable activity pattern at the topography, sources or networks level. These BS and their transitions over time reflect essential computational properties of the brain, supporting brain function and enabling cognition (Ritter et al. (2015)). The present paper focusing on solving the BS allocation problem at the network levels, that is, the detection of BSs characterised by transiently stationary large-scale causal interactions (directed information flow) between different brain ragions, or time-dependent Effective Connectivity

Several methods have been proposed to address the BS allocation problem, which can be classified into descriptive and explanatory methods. The former, use features of the measured signals to classify (allocate) time points into specific brain states. The latter are capable of generating or simulating the observed brain signals and their statistical propierties (e.g. non-stationary). The descriptive methods use time sliding windows (Hansen et al. (2015); O’Neill et al. (2017)), adaptive segmentation by grouping (Hansen et al. (2015); Khanna et al. (2015); Mheich et al. (2015)), among others (O’Neill et al. (2018); Preti et al. (2017)). These methods present various problems such as sensitivity to times windows parameters definition, instability of the estimation of the features of interest (e.g signals’ covariance matrix), among others, (for more details see (Trujillo-Barreto et al. (2019), Vidaurre et al. (2018)). The explanatory methods use generative models for characterizing the observed signals (Olier et al. (2013)). For this case, HMM has proven to be a very useful tool for solving the BS allocation problem (Baker et al. (2014),Vidaurre et al. (2018), 2016; Woolrich et al. (2013)). HMM uses a Markov chain to model the transitions between a finite set of hidden discrete BSs, such that when a BS is active it emits an observation. Although HMM addresses many of the short comings of descriptive approaches, it has important limitations as a generative model of brain activity. To resolve this limitation the use of a HsMM has been proposed (Trujillo-Barreto et al. (2019)). HSMM is a generalization of HMM where the Markov assumption leading to the geometric duration constraint is relaxed to allow for the explicit modeling of the BS duration distribution. The advantages of HSMM over HMM for the modeling of EEG amplitude fluctuations (signal envelope data) are demonstrated in (Trujillo-Barreto et al. (2019)). In this paper, a BS was characterised by a recurrent topographic (spatial) pattern of quasistatic EEG power (or brain activation)

In the context of network level BS allocation, previous work have capitalised on the flexibility and simplicity of Multivariate Autorregressive (MAR) models for modeling causal brain networks. In a MAR, the output signals represent the temporal evolution of the activity of a node (a brain area) of the brain network; and they are modeled as a linear combination of the paste activity of the other nodes. Therefore the coefficients of the linear combination (MAR coefficients) provide a characterisation of the causal interactions (directed information flow) between different nodes of the brain network (Olier et al. (2013),Vidaurre et al. (2018), Baccal á and Sameshima (2001); Kaminśki et al. (2001);Harrison et al. (2003)). Olier et al. (2013), one of the first model that use switching MARs to model time-dependent causal network, implemented a BS allocation method using a Bayesian generative model where brain signals were temporally clustered (segmented) into time periods (i.e. BS) characterised by stationary MAR models. Therefore (Olier et al., 2013) proposes adding a level of flexibility using temporal clusters characterized by MAR models. However, with this approach it is not possible to obtain a transition probability between mesostates. (Vidaurre et al., 2016) proposes HMM for model the BS transitions. This approach however, inherits the shortcomings intrinsic to HMM. In the present paper a new model that capitalises on the advantages of both HsMM and MAR is proposed to solve the brain BS allocation problems at the network level. Specifically, the HsMM for EEG proposed in (Trujillo-Barreto et al. (2019)) is adapted for the modeling of recurrent quasi-stationary large-scale causal networks in the source domain (time-depend Effective Connectivity). This is done by modeling the BS emissions as the output of a MAR that models the directed information flow (casual network) characterising the transient BS.

The identification (inversion) of the new model based on data is challenging given the high number of parameters involved. For example, if we consider that the brain consists of 62 areas (based on Desikan Killiany parcelling (Klein and Tourville, 2012)), and we use a MAR order of 10 and 10 brain states, a total of 384, 400 parameters have to be estimated (62 ∗ 62 ∗ 10 ∗ 10). Each coefficient is modelled as a mean and covariance matrix, so the actual number of parameters to be estimated is a lot more. Such a complex model can lead to overfitting or identifiability issues if enough data is not available (Bishop (2006); Ying (2019)). This problem can be addressed in different ways. In Vidaurre et al. (2018), the authors introduced a parameterisation of the MAR matrix of coefficients which reduced the number of paramaters to be estimated. This parameterisation, although mathematically convenient might impose physiological unrealistic constraints on the model. Another option is ti increase the available data to fit the model by concatenating several subject’s data along the time dimension (Vidaurre et al. (2016)). However this approach is only valid if BS features can be assume to be invariant across subjects. Another option is to pre-select a limited set of brain areas in advance in order to reduce the dimensionality of the MAR. The risk of this solution is the appearance of erroneous connections due to the interference of areas that are not included in the study (Hamilton (1994), Mardia et al. (1979)).

An alternative approach to reduce the number of parameters is use the structural connectivity information to drastically reduce the total number of possible connections and set order of the MAR based on the estimated time delays between nodes that are consistent with the structural information such as fiber lengths (Fukushima et al. (2015),Sporns et al. (2000),Stephan et al. (2000)). That is, functional integration between different areas in the brain are mediated by white matter (WM) connection (Sotiropoulos and Zalesky, 2019). Studies have reported that functional dynamics are a reflection of structural connectivity (Damoiseaux and Greicius, 2009). (Koch et al., 2002) so that time-depend connectivity fluctuate around a structural connectome ((Nakagawa et al., 2013; Deco et al., 2017; Cabral et al., 2017)). These findings suggest that anatomical connectivity can be used as a constraint in the study of functional connectivity. This type of constraint have been used in previous models to estimate static functional connectivity ((Penny et al., 2005), (Fukushima et al., 2013), (Fukushima et al., 2015)). In the present paper, we evaluate the benefits of using structural connectivity constraints in the estimation of tdEC. Although structural information in this work was derived from the human connectome, the proposed approach could also be used to incorporate individual structural information should sush data is available.

In studies such as (Penny et al., 2005), (Fukushima et al., 2013) and (Fukushima et al., 2015) structural connectivity is applied as a constraint to MAR models of high complexity. However, these studies do not take into account the dynamics of brain states. In addition, an arbitrary threshold is applied in (Fukushima et al., 2015) to define the structural constraint.

## 2 METHODS

We describe a HSMM-MAR-AC model for BS allocation, where BS is defined as a specific configuration of an effective brain network, that persists over short-periods of time, generating a continuous sequence of the observed brain activity, before switching to another BS. We elucidate how the connectome information can reduce the description of the model, and hence the number of parameters to be estimated. We describe the inversion procedure to estimate the parameters using the Bayesian machinery, and the performance metrics and experiments used to evaluate the accuracy and efficiency of the model.

### 2.1 The Generative Model

#### 2.1.1 MAR Model Description

In the MAR model, the neuronal activity of each area is characterized as a node of the network. The activity of a node X corresponds to the linear combination of the activity in the past (with delay) of the other areas connected to X. The interaction between the nodes is not instantaneous, each interaction will take place in a specific delay (1 to L). Furthermore, each interaction towards X will be modulated by an Autoregressive coefficient. Finally, a Gaussian noise is incorporated into the resulting neuronal activity. This can be represented as follows:

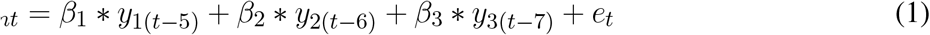

Where *y*_*nt*_ correspond to the neuronal activity of the nodes n at time *t, β*_*n*_ denote the autoregressive coefficient and *e*_*t*_ is the noise Gaussian. In this case there is a neural connection from node 1, 2 and 3 to node n which implies that the neuronal activity of node n is determined by the linear combination of the activity of nodes 1, 2 and 3 at delays 5, 6 and 7 respectively.

#### 2.1.2 HsMM-MAR-AC Model Description

The model comprises two components: The HsMM characterizes the time-dependent brain effective connectivity as rapid transitions between discrete BSs that persist over short duration in time Trujillo-Barreto et al. (2019). For each state, the brain activity in each brain area is defined by the activity and effective (time-lagged) connections among brain areas, described by the second component, the MAR, forming a stationary (or quasi-stationary) network (Penny and Roberts (2002),Olier et al. (2013),Vidaurre et al. (2016)). Figure 1 shows the dynamic Bayesian network (DBN) graph representation of the model, together with the conditional dependencies between the parameters, hidden variables, and the observed data.

**Figure 1.**
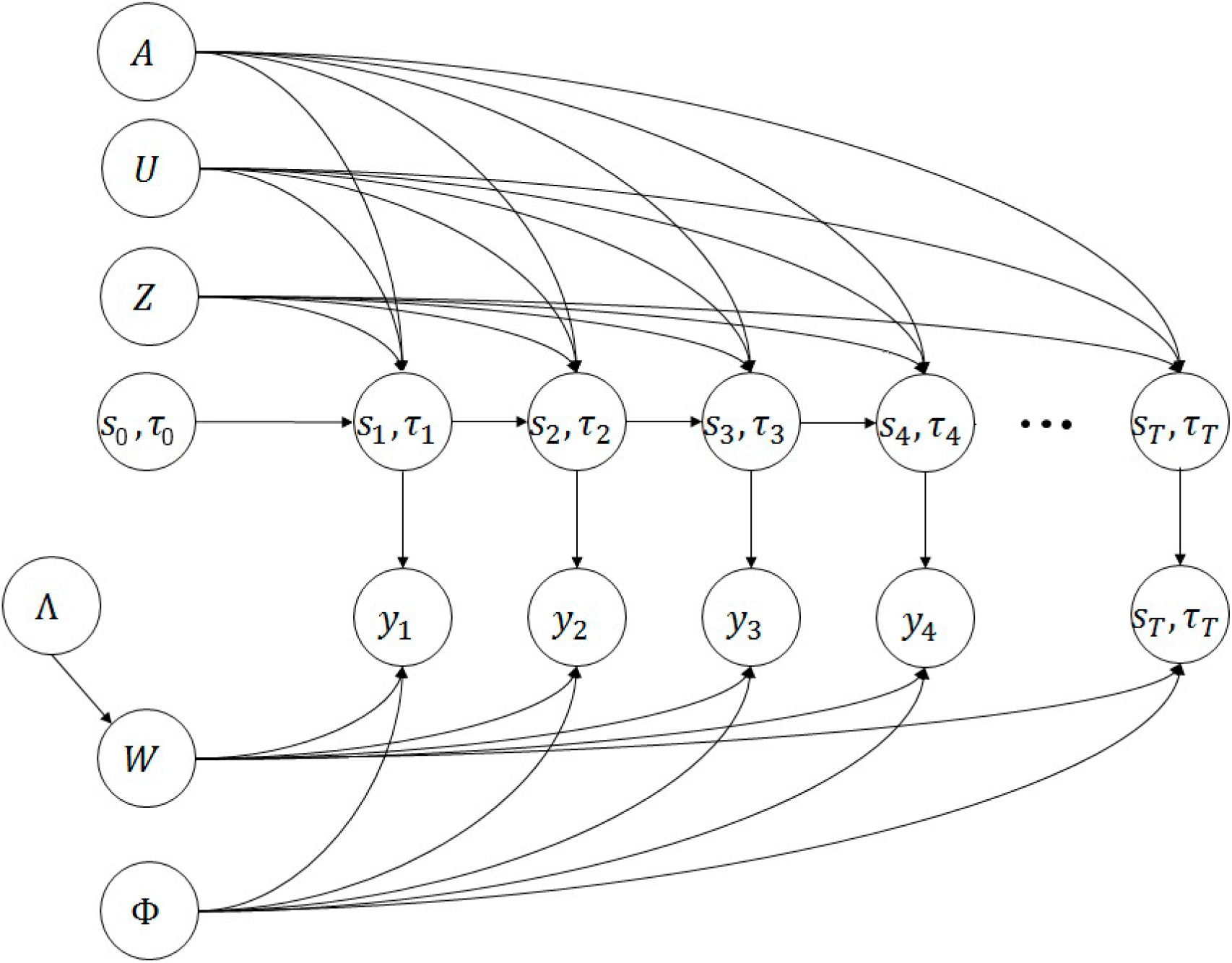
Dynamic Bayesian graph of the generative probabilistic HsMM-MAR model. When in state *s*_*T*_ the model emits a sequence of measurable brain activity *y*_*T*_ according to the rules specified by the MAR coefficients in *W* ^(*M*)^, for a duration given by *U* = *µ*^(*M*)^ and *Z* = *ζ*^(*M*)^. When *s*_*t*_ runs out of time *τ*_*t*_, the model switches probabilistically to another BS according to the transition matrix *A*.

**Figure 2.**
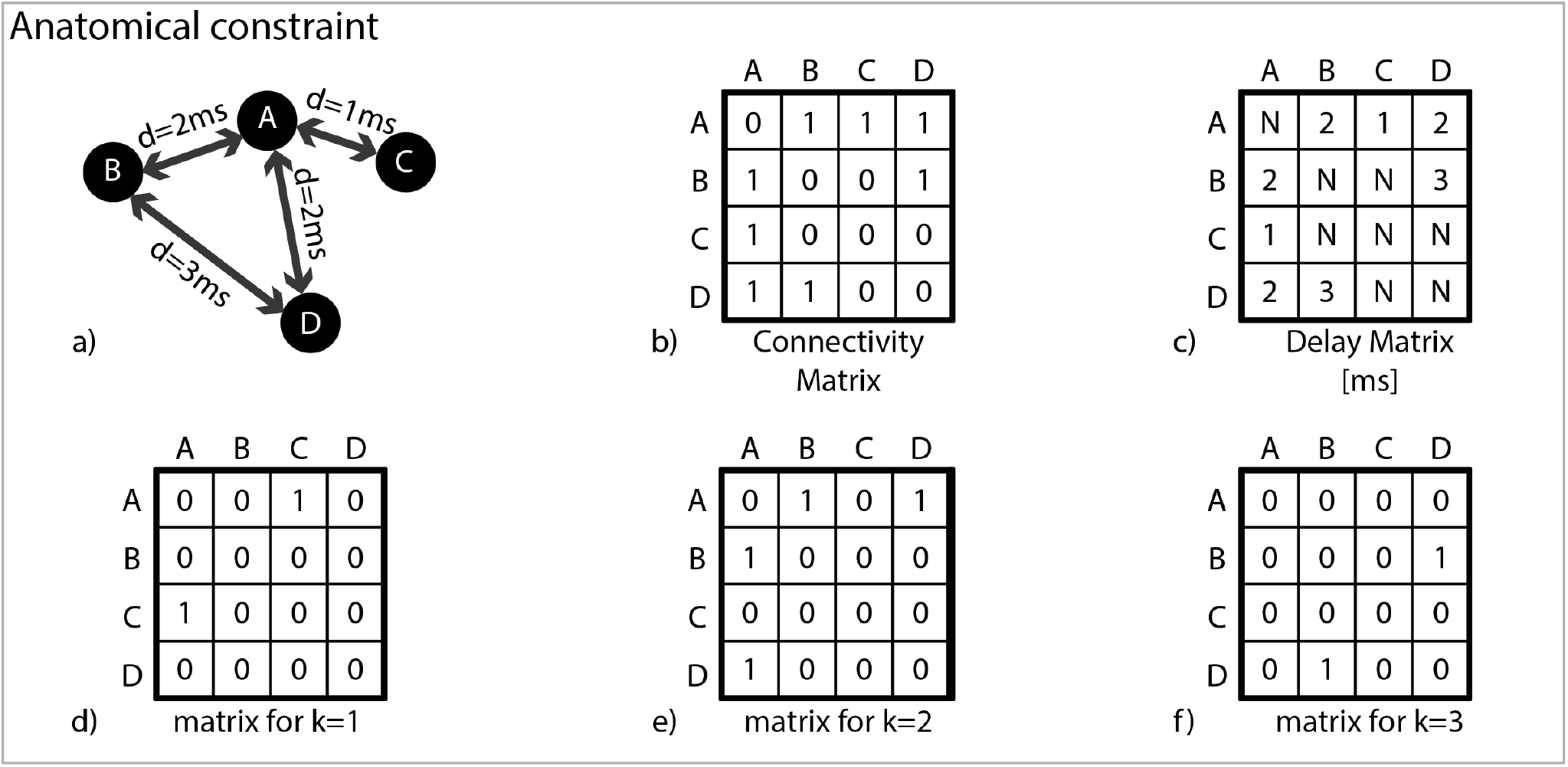
Using Anatomical Connectivity information to constrain the MAR emission model. a) In this schematic example there are 4 brain areas, A, B, C and D. In the absence of any constraints, and assuming a maximum AR lag of 5, there would be 4 ∗ 4 ∗ 5 = 80 parameters to estimate, for each BS. The model is simplified in two ways: b) Applying Structural Connectivity (see section 2.2.1), MAR coefficients where there no are anatomical connections, are dropped; and c) for each anatomical connection, one and only one AR lag is considered. That lag is determined based on the transmission delay in that connection, which in turn is calculated from the connection length, see section 2.2.2. In this example, there are only 4 connections with delays between 1 and 3 ms, and for simplicity we assume each MAR lag corresponds to 1ms. d-f) The MAR coefficients reduce to 4 ∗ 4 ∗ 3 matrices, with a total of 8 parameters to be estimated. The connections and delays are symmetric, but coefficients are not.

Where:

N: Number of nodes M: Number of states L: Maximun delay

*y*_*t*_ : Observed current density (brain activity) at time *t* in each brain area.

*s*_*t*_ : Hidden BS at time *t*. Here there are M states.

*τ*_*t*_ : Remaining time in the BS *s*_*t*_.

*W* : Set of matrices of MAR coefficients, one for each BS, *W* = {*W* ^(1)^, *W* ^(2)^, …, …*W* ^(*M*)^} and Λ set of matrices that contain their corresponding precisions Λ = {Λ^(1)^, Λ^(2)^, …, …, Λ^(*M*)^}

Φ : Set of vectors of observation noise precision.Φ = {*ϕ*^(1)^, *ϕ*^(2)^, …, …, *ϕ*^(*M*)^}. *A* : M x M Transition probabilities matrix between BSs.

*U* : Set of M values that denote the mean duration of each BS, *U* = {*µ*^(1)^, *µ*^(2)^, …, …, *µ*^(*M*)^} and *Z* : is a set of M values that denote their corresponding precisions *Z* = {*ζ*^(1)^, *ζ*^(2)^, …, …, *ζ*^(*M*)^}.

The full likelihood function of figure 1 is expressed as:

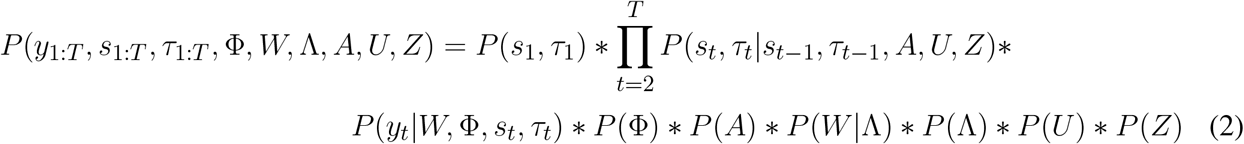

The probability of emitting an observation *y*_*t*_ at instant *t* given that the state *s*_*t*_ of duration *τ*_*t*_ corresponde to:

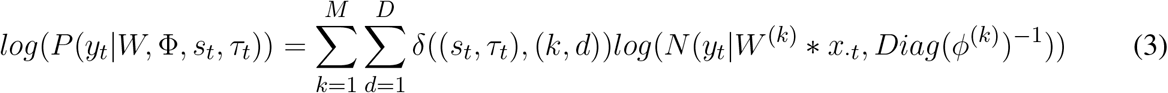

Where, *x*_*·t*_ = (*y·*_(*t−*1)^*T*^_, *y*_*·*(*t−*2)^*T*^_,,, *y*_*·*(*t−l*)^*T*^_), denotes the *t* column of the matrix *X* corresponding the past observations in each brain area, i.e. the regressors. The number of autoregressive coefficients to estimate in equation 3 are the elements of matrix W, which corresponds to a full matrix of dimensions N * N * L.

For example, for a brain defined by the DKT atlas (Klein and Tourville, 2012), which contains 62 areas, a maximum lag of 10, and 9 brain 7states, the number of autoregressive coefficients to estimate is 62*x*62*x*10*x*7 = 269, 080.

Here we use anatomical connectivity information to reduce this number in two ways: Limiting the connectivity matrix to only where structural connectivity exists (see section 2.3.1), and use the tractography data to calculate the precise fibre length thus estimate exact delay between two regions (section 2.3.2). The exact delay allows restricting the number of lags per connection to just one. In the above example, if half the connections were considered and one lag is estimated, the number of coefficients drops to 1/2 × 62 × 62 × 1 × 7=13,454, which is around to 6% of the original number of coefficients.

To implement this reduction, the multivariate Gaussian probability density of the N areas in equation 3 is separated into N uni-variable Gaussian distributions. This is feasible since the precision matrix *Diag*(*ϕ*^(*k*)^) has only elements on the diagonal (they are considered independent). For source n we have that the new precision is 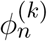 and corresponds to element n of the diagonal of the matrix *Diag*(*ϕ*^(*k*)^).

In addition, the Gaussian mean described in equation 3 by *W* ^(*k*)^ *∗ x*_*·t*_ is modified by 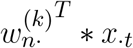 where 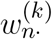 denotes the vector of column n of the coefficient matrix W. This is where the anatomical information is used, which is translated in having a vector *l*_*n*_ extracted from the connectivity matrix estimated in 2.2.1 that contains the indices of those connections not discarded for source n. That is, those connections that are most likely and that correspond to the estimated delay. Using this vector we have the mean of the Gaussian of the source n we can write it as 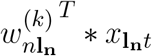 where these variables correspond to a subset of 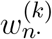 and *x*_*·t*_ respectively, based on the indices of *l*_*n*_. With this modification, equation 3 becomes equation 4.

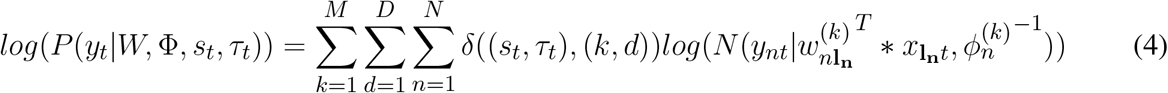

#### 2.1.3 Priors

We model the MAR coefficients using a zero mean multivariate Gaussian and precision matrix with only elements on the diagonal, which considered independence among the coefficients (equation 5).

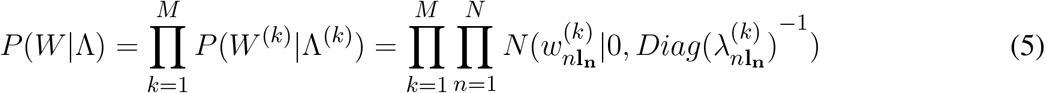

Where 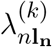 corresponded to a subset of 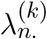 according of index of *l*_*n*_ in state k.

The Gaussian distribution dimension that describe 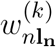 corresponded to the total number of indices contained in *l*_*n*_.

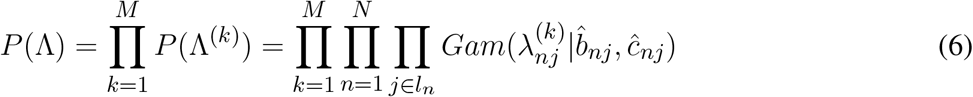

Where *λ*_*nj*_ corresponded to the j-component of the vector 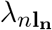. In addition, 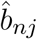 and 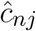 corresponded to the shape and scale parameters of the Gamma distribution. Their values are 1000 and 0.001 respectively.

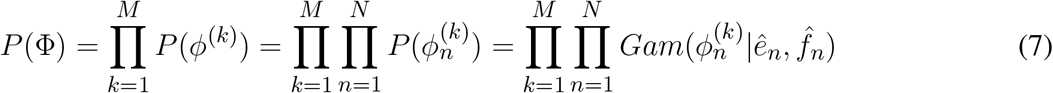

Where ê and 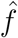corresponded to the shape and scale parameters of the Gamma distribution. Their values are 1000 and 0.001 respectively.

For the description of the other model parameters (transition, duration of the states and initial state) see Trujillo-Barreto et al. (2019)

#### 2.1.4 Model Parameters Estimation

No analytical solution were found in the parameters model estimation; thus, the parameters estimation was performed using an approximate inference method called Bayes Variational (Bishop, 2006) and Beal (2003). VB used an auxiliary function *q*(*x*) to approximate as closely as possible the posterior distribution of the parameters and/or hidden variables. Mean Field Approximation Jordan et al. (1999) allowed to chose a factorizable *q*(*x*). Conjugate priors obtained the analytical expression of updated the parameters and hidden variables, in time. The mean field approximation used for the joint probability was:

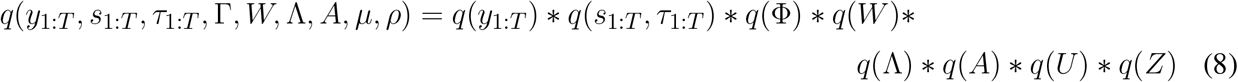

For full details about model parameters and their estimation, see supplementary material.

### 2.2. Anatomical Constraint

#### 2.2.1 Structural Connectome Calculation

Whole-brain probabilistic tractography was reconstructed using the software MRtrix, based on spherical deconvolution model of (Tournier et al., 2007) and dMRI data of 106 subjects from the HCP database (http://www.humanconnectomeproject.org). Anatomically-Constrained Tractography (ACT) (Smith et al.,2012) was calculated with: step size = 0.1 *∗ voxelsize*, angular threshold= 90, minimum length = 30*mm*, maximum length = 250*mm*, cutoffvalue = 0.06, and 30 million strles. Each tractography dataset was filtered with SIFT tool (Spherical-deconvolution Informed Filtering of Tractograms, (Smith et al., 2013), maintain 3 million of fibers. The tractography of each subject was normalized to the MNI space using the corresponding non-linear transformation.

Figure 3a) shows the pipeline used for calculating the average weighted matrix. We computed the connectivity matrix with the probabilistic tractography and the anatomical Desikan Killiany (DKT) atlas (Klein and Tourville, 2012). This matrix contains the numbers of fibers (*F*_*ij*_) connecting ROI_*i*_ and ROI_*j*_. Figure 3b) illustrates the approach for calculating the anatomical connectivity. We chose the most repeated ROIs evaluating a neighborhood of 5*mm* of each extreme point. This approach, combined with ACT, obtains better labelled than when using the extreme point, or all the voxels intersecting the path of the fiber (Yeh et al., 2019). Hence, we constructed the individual connectivity matrix (62 ∗ 62) representing the *F*_*ij*_ of each pair of anatomical ROIs. We calculated the individual weighted connectivity matrix (*w*_*subjecti*_), based on the connection strength (*w*) between each pair of ROIs (equation9 (Cheng et al., 2012; Dimitriadis et al., 2017)). Where *n*_*i*_ and *n*_*j*_ were the volume of ROI_*i*_ and ROI_*j*_, respectively. The average weighted connectivity matrix *w*_*mean*_ was calculated, from the 106 individual weighted connectivity matrices

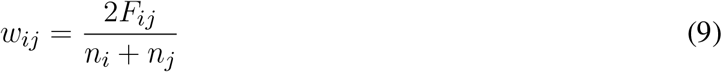

We used *w*_*mean*_ and thresholds for defining the percentage of connections, which we used to define different structural constraints including the reference structural constraint, see 2.2.4.

**Figure 3.**
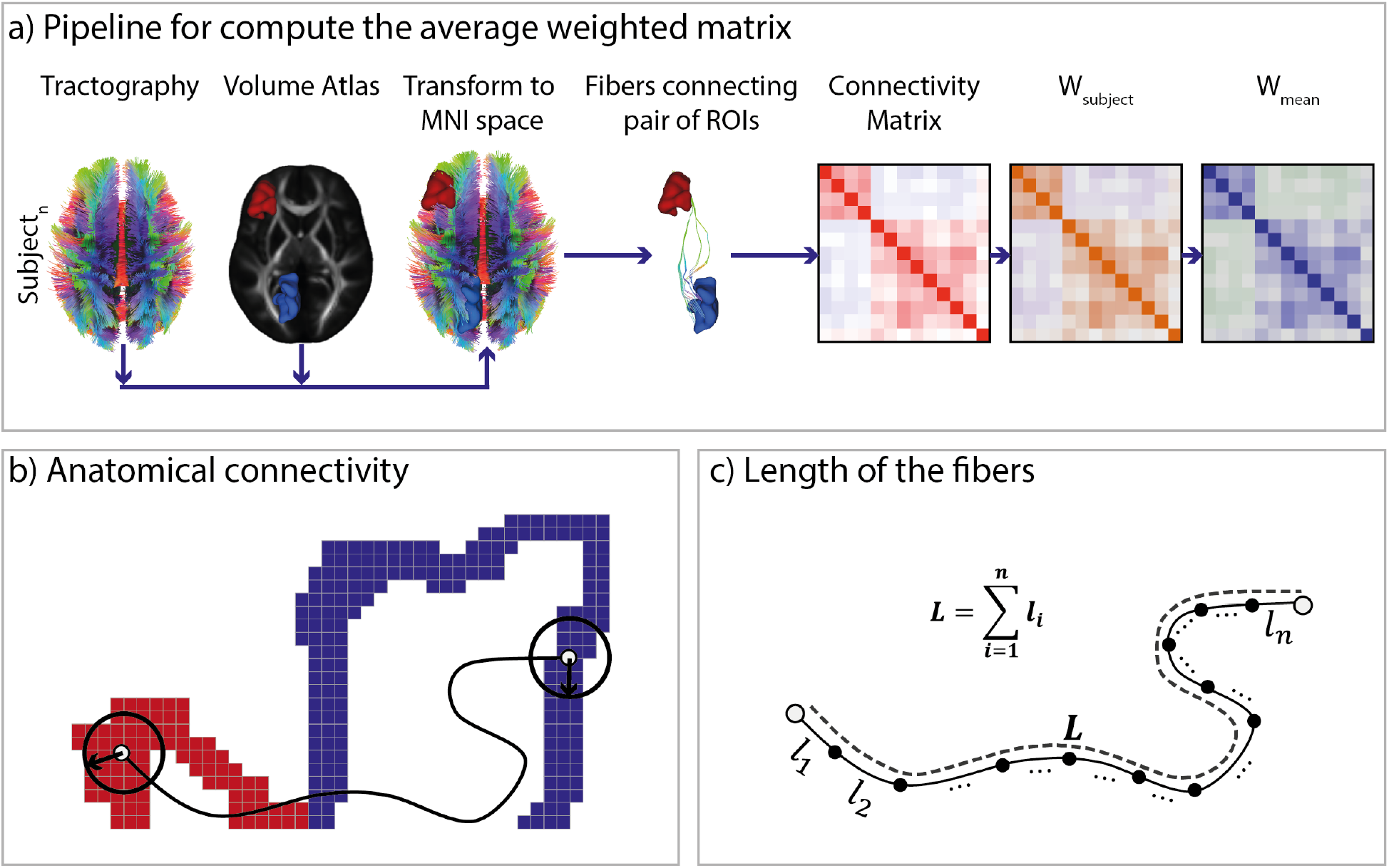
a) Calculation of the average weighted matrix *w*_*mean*_. Subject connectivity matrices are first calculated, based on the tractography and anatomical ROIs in MNI space. Then, individual weighted connectivity matrices (*w*_*subjecti*_) are computed based on the calculation of the connection strength (w) between each pair of ROIs. Finally, *w*_*mean*_ matrix is computed as the average of each matrix. b) Method for calculating anatomical connectivity using a radius at the ends of the fiber. c) Method for calculating the length of each fiber.

#### 2.2.2 Calculation of the MAR lags

We calculate the conduction delay between two connected regions as the average path length of connecting fibres divided by the conduction velocity for myelinized axons of 6 *m/s* (Ghosh et al., 2008), and 3 *m/s* for self-connections (Fukushima et al., 2015). The path length of each fiber (*L*) was calculated by adding the Euclidean distance between consecutive points, figure 3c). The conduction delay is converted to MAR lag by dividing the delay by the temporal resolution of the data, here 5ms.

#### 2.2.3 Resting state networks

We used the seven most reported RSNs. Default Mode Network (DMN) (Corbetta and Shulman, 2002; Greicius et al., 2003; Fox et al., 2005; Raichle and Snyder, 2007; Finn et al., 2015; Buckner and Dinicola, 2019), Sensori-motor Network (SMN) (De Luca et al., 2005; Fox et al., 2006; Finn et al., 2015), Executive Control Network (ECN) (Seeley et al., 2007), Visual Network (VN) (Himberg et al., 2004; Abou Elseoud et al., 2009; Hasson et al., 2009; Stevens et al., 2010; Finn et al., 2015), Fronto-Parietal Network (FPN) (Fox et al., 2005; Damoiseaux et al., 2006; De Luca et al., 2006; Smith et al., 2009; Albert et al., 2009; Finn et al., 2015), Auditive Network (AN) (Cordes et al., 2000; Kiviniemi et al., 2003; Van Den Heuvel et al., 2008) and Temporo-Parietal Network (TPN) (Kiviniemi et al., 2003; Dronkers et al., 2004; Van Den Heuvel et al., 2008). Regions of networks were defined according to the most reported cases, and finding their correspondence with the ROIs of DKT atlas

#### 2.2.4 Anatomical constraint

We applied a threshold to *w*_*mean*_ which guaranteed that each node of the seven networks was connected, regardless the self-connection. Under this condition, we obtained a matrix with 28% of connections, which with the delay matrix were used to constructed the reference structural constraint. In addition, structural connectivity matrices were constructed with different percentage of connections (15%, 20%, 35%, 40% and 100%).

### 2.3 Simulations

Two experiments were performed, the first aimed to verify the performance of HsMM-MAR-AC using synthetic data generated from a small generic system. The second consisted of testing the HSMM-MAR-AC model using synthetic data as close as possible to the real functional data in terms of magnitude and composition of the networks. It consisted of the 10 non-stationary signals composed of the 7 most reported RSN in the literature. This simulation aimed to verify the limitations and advantages of the model in a real usage scenario. The methodology is explained in more detail in the following sections.

#### 2.3.1 Experiment 1 (Model Validation)

The first simulation consisted of generating data using a 10-node and generative HSMM-MAR-AC with a maximum delay of 3. Simulations are performed using 3, 4 and 5 states and the network characterizing each state uses 6 of the 10 nodes. Theses MARs are created randomly under condition that they are stable according to the procedure explained in (Gambetti, 2011). We used a single delay for connection among nodes (there were no multiple connections). In addition, we implemented a random ranking to provide a fixed score for the connections, then the connection with lower scores were discarded. Thus, a degree of structural constraint was defined to discards connections based on the ranking. The data was performed using a 50% of structural constraint, i.e. with 50 of 100 possible connections (10 * 10 nodes). Then we use the generated data to train a HSMM-MAR-AC model which is evaluated according to 2.4. The model was trained using different structural constraint (25%, 50%, 75% and 100%) adn different data length. For each evaluation, 10 runs are executed with different MAR conformations.

#### 2.3.2 Experiment 2 (Simulation and Estimation of RSN)

Figure 4b) shows the approach used to generate the signals. Ten non-stationary signals were generated based on the reference connectivity (28%), which included the seven most reported RSNs, refer section 2.2.3. HsMM generative model (Trujillo-Barreto et al., 2019) was used to generate the signals. Each network was represented by a MAR (*A*_*i*_), where *i* is the *i*_*th*_ RSN. Each network *i*, had a *S*_*i*28%_ matrix, which contains only the lags for the regions in network *i*. In addition each network had a stable MAR process Ã_*i*_ with all ROIs connected for all lags. A MAR process is stable when the module of all its eigenvalues is inferior to 1.0 (Gambetti, 2011). The final MAR of each RSN (*A*_*i*_) were obtained using the equation 10, which the operator ⊙ means point to point multiplication. Longer average fiber was 239.8*mm*, thus the maximum connection delay was 40*ms*, and the maximum lag was k=8, considering a velocity of conduction of 6*m/s* and a temporal resolution of the signal _*t*_ = 5*ms* (see section sec: 2.2.4). The DKT atlas has 62 ROIs, thus, the dimensions for *S*_*i*28%_ was (62 ∗ 62 ∗ 8). The probability distribution of state duration was a log-normal constructed with a mean of 3.84 and 0.4% of standard deviation. The transition probability matrix was designed with a non-symmetric structure, avoiding to favor particular transition paths.

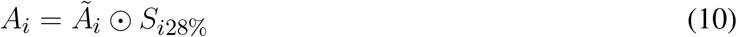

Figure 4c) illustrates the estimation stage, which the signals generated in the simulation stage were used to estimate the model parameters. We used separately six different anatomical constraints, (refer to struct-matrix). We increased the size of the estimated signals in steps of 120*s*, from 240*s* to 600*s*. The initialization of the algorithm were random. The model estimated the number of states, the state sequence, the transition probability matrix and the auto-regressors of MAR states (*A*_*i*_).

**Figure 4.**
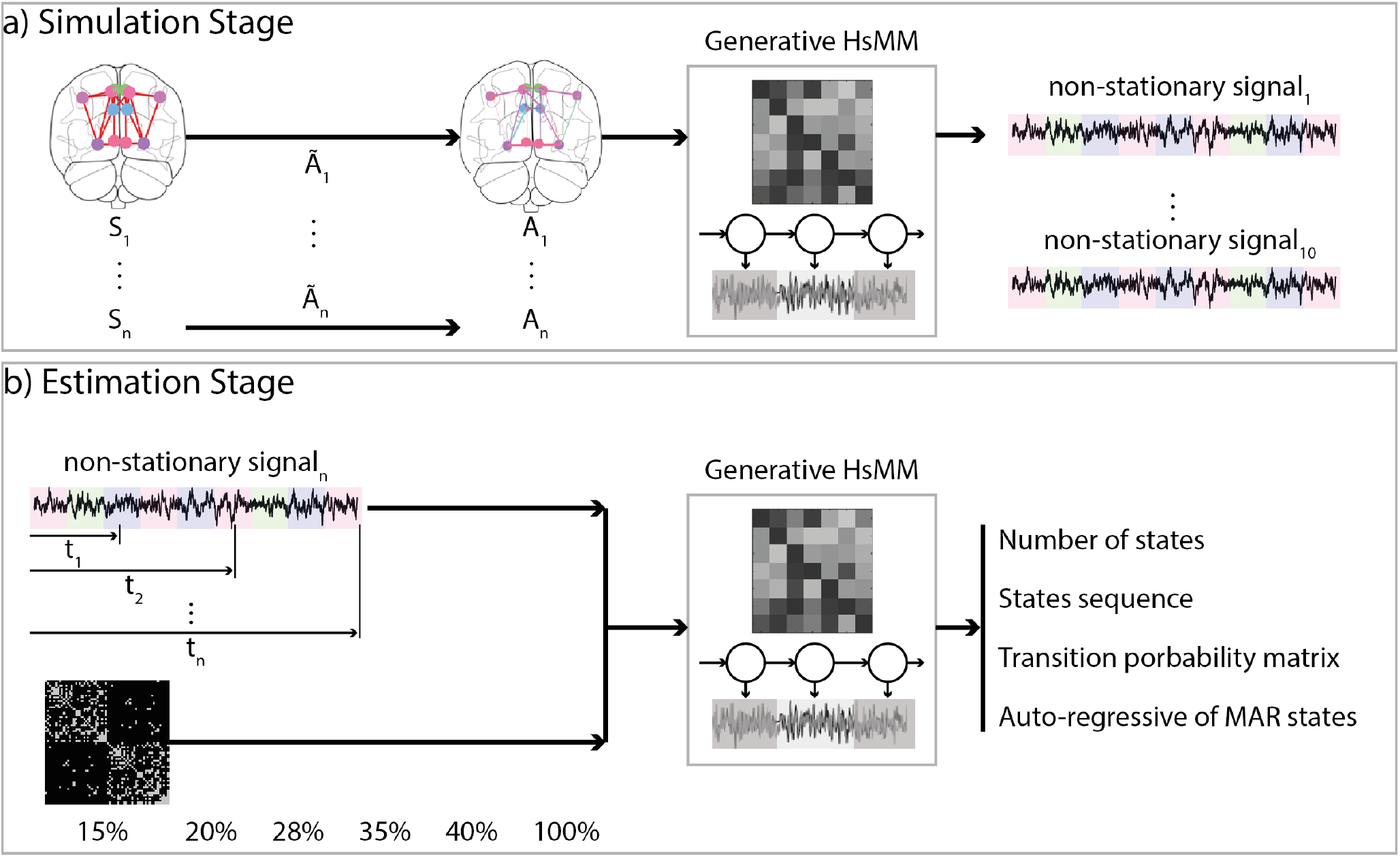
Figure shows the general scheme of simulation and estimation in a) and b), respectively. a) Simulation stage, we construct the structure (*S*_*i*_) of 7 most reporter RSNs and we generate 7 stable-MAR process Ã_*i*_. Then, we multiply them point by point and we obtain the *A*_*i*_ for each network. In addition, we use the generative HsMM-MAR for generating 10 non-stationary signals. b) Estimation stage, we use the 10 non-stationary signals with the structural constraints. Then, we use the HsMM-MAR for estimating the number of states, the states sequence, the transition probability matrix and the auto-regressive of MAR states.

**Figure 5.**
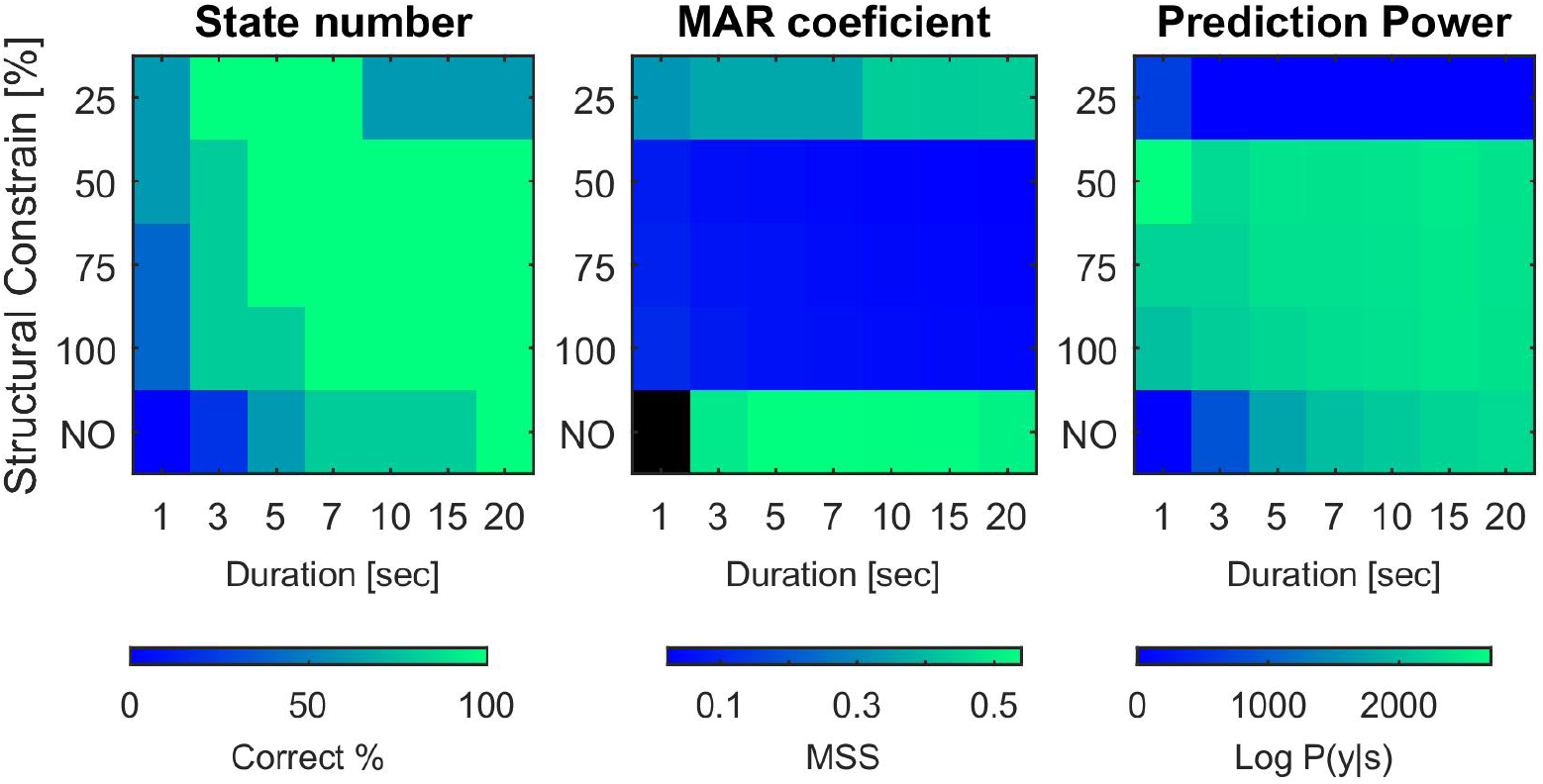
In the three graphs, the Y axis indicates the % of AC used and the X axis the duration in second. Figure A shows the % recovery of the number of states over a total of 10 runs. As can be seen, with a AC of 50% and a data length of 5 sec, convergence is achieved 10 times. In the case that a AC of 100% is used (only the delay is restricted) a data length of 7 sec is required to converge correctly the total number of times. Only when AC is not used, that is, connections and delay are not restricted, does the requirement grow, reaching 20k. In the case of not using AC and using less than 5 sec, the probability that it converges correctly is almost zero. Figure B shows the recovery performance of the MAR coefficient matrix by means of the MSS distance between the real matrix and the estimated one. The range of this metric is from 0 to 1 and it was obtained only for those cases where the number of states was correctly determined. The black color indicates that it is not possible to calculate this metric because there are no cases where it converges to the correct number of states. In most cases the recovery is very good, resulting in values around 0.1. Only in those cases of not using AC or using 25%, the recovery performance degrades. Figure C shows the predictive power of the model. Only values between the same column are comparable. Between different columns are not comparable since the data is different. It can be seen how the maximum prediction power is reached for 50% of SC. The worst cases are with 25% AC and without SC. It can be seen that as the AC increases, the prediction power decreases slightly, with the maximum always in AC being 50%.

**Figure 6.**
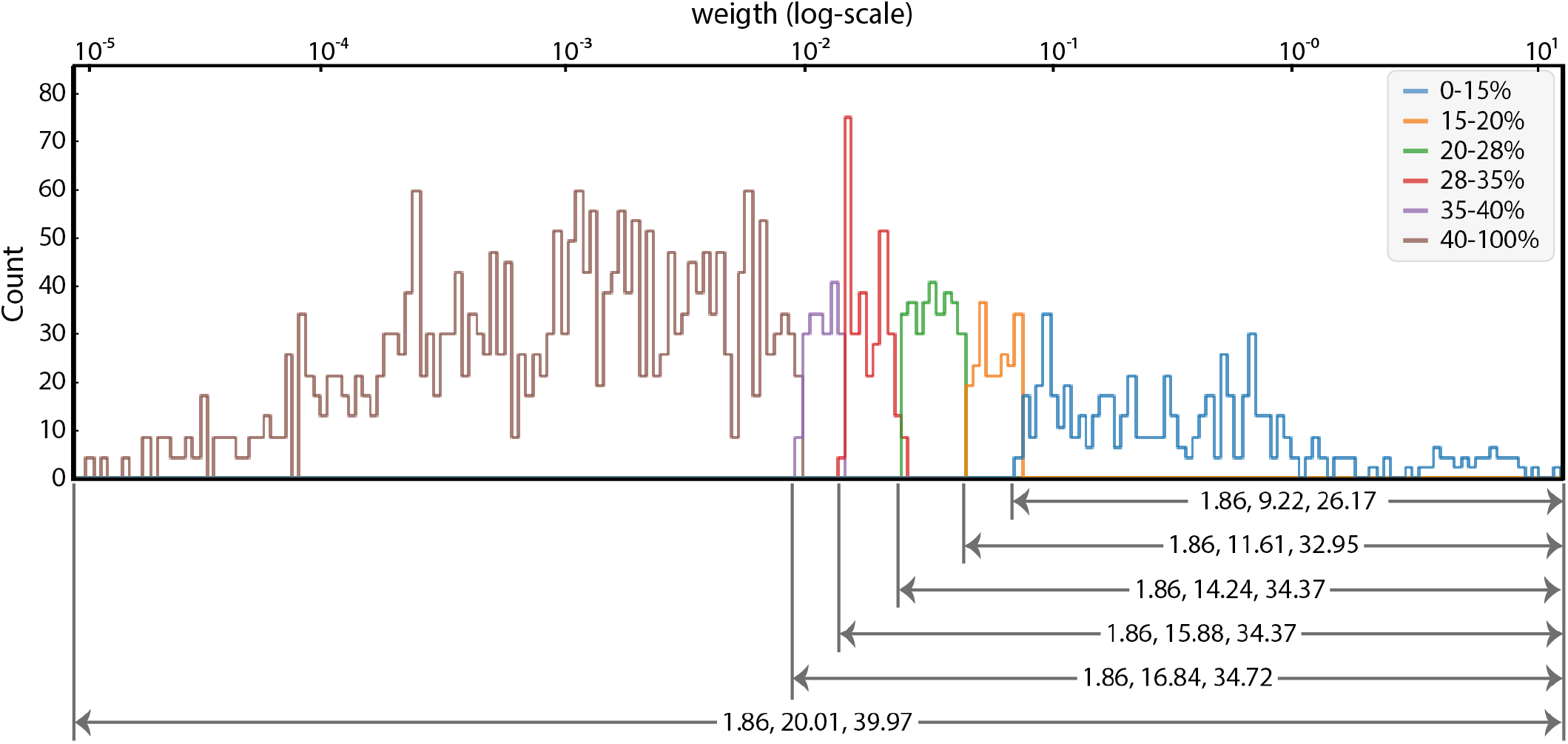
a) Histogram of the connection weight. The colors show the histogram of the different structural constraints. b) Minimum, mean and maximum value of the connection delay (in millimeters) corresponding to each structural constraint. With stronger restrictions, some long connections were lost. However, with the 15% structural constraint, only short ECN and DMN connections were lost.

### 2.4 Model Evaluation

#### 2.4.1 Similarity matrix

MSS similarity matrix (Sotero et al., 2010) was used to evaluated both the performance in estimating the MAR of each state and the transition matrix. This metric was defined in equation 11, which *M* is the simulated matrix and 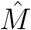the estimated one. The values ranged from 0 to 1, being 1 for identical matrices.

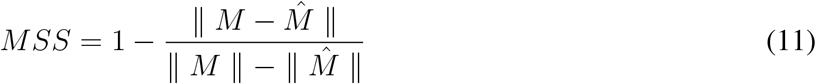

#### 2.4.2 Kullback–Leibler divergence

Analytical Kullback–Leibler (KL) (Joyce, 2011) divergence was used to evaluated the performance of the algorithm for recovering the duration distribution (equation 12).

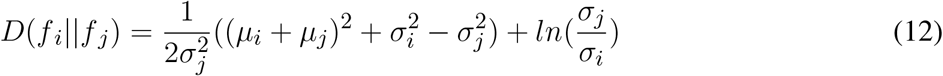

#### 2.4.3 G-test

G-test (Hoey, 2012) was used to determine if the estimated duration parameters significantly differed from the design parameters (equation 13).

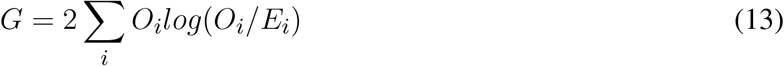

where *O*_*i*_ and *E*_*i*_ were the observed and expected values, respectively.

## 3 RESULTS

### 3.1 Experiment 1: Model Validation

In this simulation, the performance of the HSMM-MAR model is evaluated using simulated data from a small network (10 nodes), maximum lag of three and an structural constraint (AC) of 50%. Performance evaluation is performed based on the length of the training data sequence and the AC used in model training. In addition, the evaluation of the model is carried out in two aspects: the verification of the recovery of the parameters of the original model and the evaluation of the prediction capacity of the model in new data. For more details on the evaluation see 2.4.

The first parameter to be evaluated is of the structural type and corresponds to the number of states. This parameter is very important, since its incorrect estimation involves the impossibility of recovering the other parameters of the model. For this evaluation, data with 3, 4 and 5 states (networks) and with a data sequence length of 5 second (5 thousand point) and 20 second are generated. Then, based on the data, the models are trained and the convergence curves of the selection of the models are obtained, which allows us to estimate the number of states. The result of the convergence curves can be seen in Supplementary Material. Here shows how the HSMM-MAR model with RE correctly estimates the number of states for all cases. On the other hand, without RE, it correctly estimates the number of states only for the case of 20 second.

Two very important variables to evaluate the performance of the models are the % of AC and the length of the data sequences used in the estimation of the model parameters. For this reason, we will evaluate the recovery of these parameters based on these two variables using a matrix where the color will indicate the performance of the model. The first thing is to evaluate the performance of the recovery of states, which can be seen in figure A of 5. In Addition, the recovery of the MAR coefficient matrix can be seen in the figure B of 5. In these figures it can be seen how the use of AC allows the model to present a good performance with a much shorter data length than the case of not using AC. This is consistent with the fact that using a CS implies the need to estimate a greater number of parameters and, therefore, a greater amount of data is required.

With respect to the recovery of the parameters associated with the transition matrix, duration distribution of the models, a good performance is observed, with no differences between the different cases. To see the details of this performance see supplementary material.

In this section, the predictive power of the model is quantified. For more details on obtaining this metric, see 2.4. These results can be seen in Figure C of 5. It can be seen how the use of AC allows reaching values close to the maximum of prediction power with a smaller data length requirement. It is also verified that the variations of AC between 50% and 100% do not significantly vary the prediction power.

Finally, regarding the recovery of the sequence of states of new data, it is verified that the model presents good performance without showing greater variation with respect to the % of AC used and the length of the data sequence. These results can be seen in Supplementary Material.

### 3.2 Experiment 2: Simulation and Estimation of RSN

We used structural constraints with different percentages of connections (15%, 20%, 28%, 35%, 40% and 100%) with 10 no-stationary signals and 5 iterations for each run, for testing the performance of HsMM-MAR. The signals were obtained using the generative HsMM-MAR with the seven most reported RSNs structure (refer section 2.2.3) and length of 240*s*, 360*s*, 480*s* and 600*s*. The reference structural constraint (28%) was selected for constructed the RSNs structure, which was found for assuming that each node of each network must be connected to, at least, one node. We used a non-symmetric transition matrix, which probabilities avoided a preferent path. The probability duration of states was designed with a mean of 3.84 and 0.4% of standard deviation. The reference structural constraint contained 1066 connections. Under this conditions the RSNs structure had the following number of connections: DMN (66), SMN (32), ECN (68), AN (34), VN (16), FPN (88), TPN (104). The structural constraints with 15%, 20%, 35%, 40% and 100% had 578, 770, 1346, 1540 and 3844 connections, respectively. The ECN and DMN networks lost connection for structural constraints of 15% and 20%. Figure 7 shows the connections for 15, 20 and 28% of connections. For 15% of connections the DMN had 44 of 66 connections and ECN had 44 of 68 connections. For 15% of connections the DMN had 50 of 66 connections and ECN had 52 of 68 connections.

**Figure 7.**
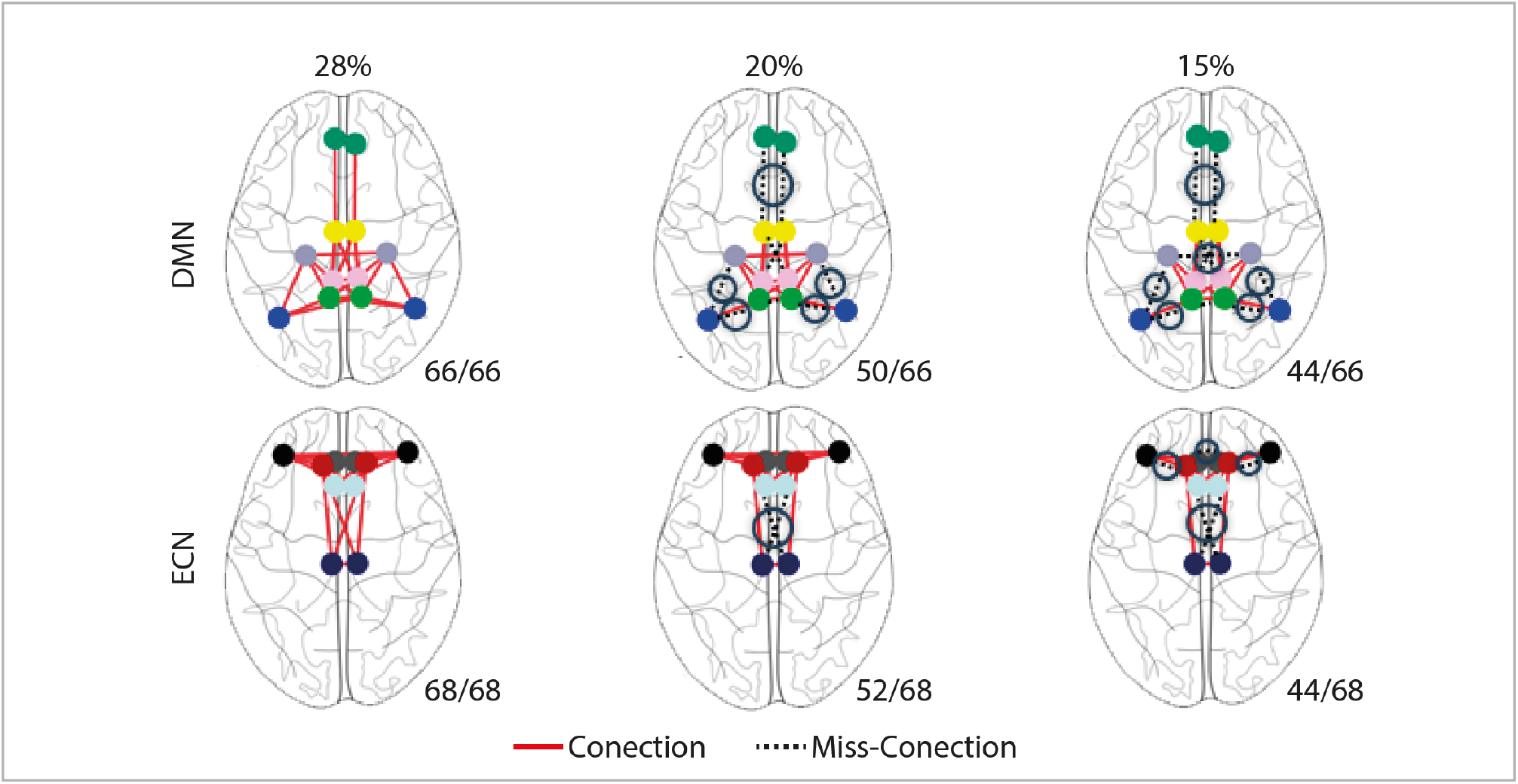
Illustration of the anatomical connections in the DMN and ECN designed for 15%, 20%, and 28% structural constraint. For each case, the lower right corner shows the percentage of connections with respect to the reference connectivity (28%). Note the loss of connections for 15 and 20% of structural constraints (in dashed lines).

The correlation between the delay and the weighted connectivity is *−*0.30 with p-value *<* 0.01. Figure 6a) shows the histogram of the connection weight. Figure 6b) shows the minimum, mean and maximum value of the connection delay for the structural constraints. Stronger restrictions eliminate long connections, which do not influence the ROIs of the RSNs.

Figure 8a) shows the percentage for detecting, correctly, the number of RSNs. HsMM-MAR was able to detect the correct number of networks with structural constraints of 15 and 20% of connections. For structural constraints larger than 20%, HsMM-MAR needed signals with, at least, 360 seconds. Higher accuracy was obtained for larger signals and smaller connections in the structural constraints. Figure 8b) shows the MSS metrics of the estimated versus simulated networks. The NA values correspond to cases where the correct number of networks was not found. Networks were matrices which represented a MAR process. Higher MSS values were found for the reference structural constraint (28%) and larger signals. Both DMN and ECN networks had smaller MSS for structural constraints of 15 and 20% of connections. Figure 8d) shows the MSS metrics of the estimated versus simulated transition probability matrix. The NA values correspond to cases where the correct number of networks was not found. Higher metrics were obtained for larger signals.

**Figure 8.**
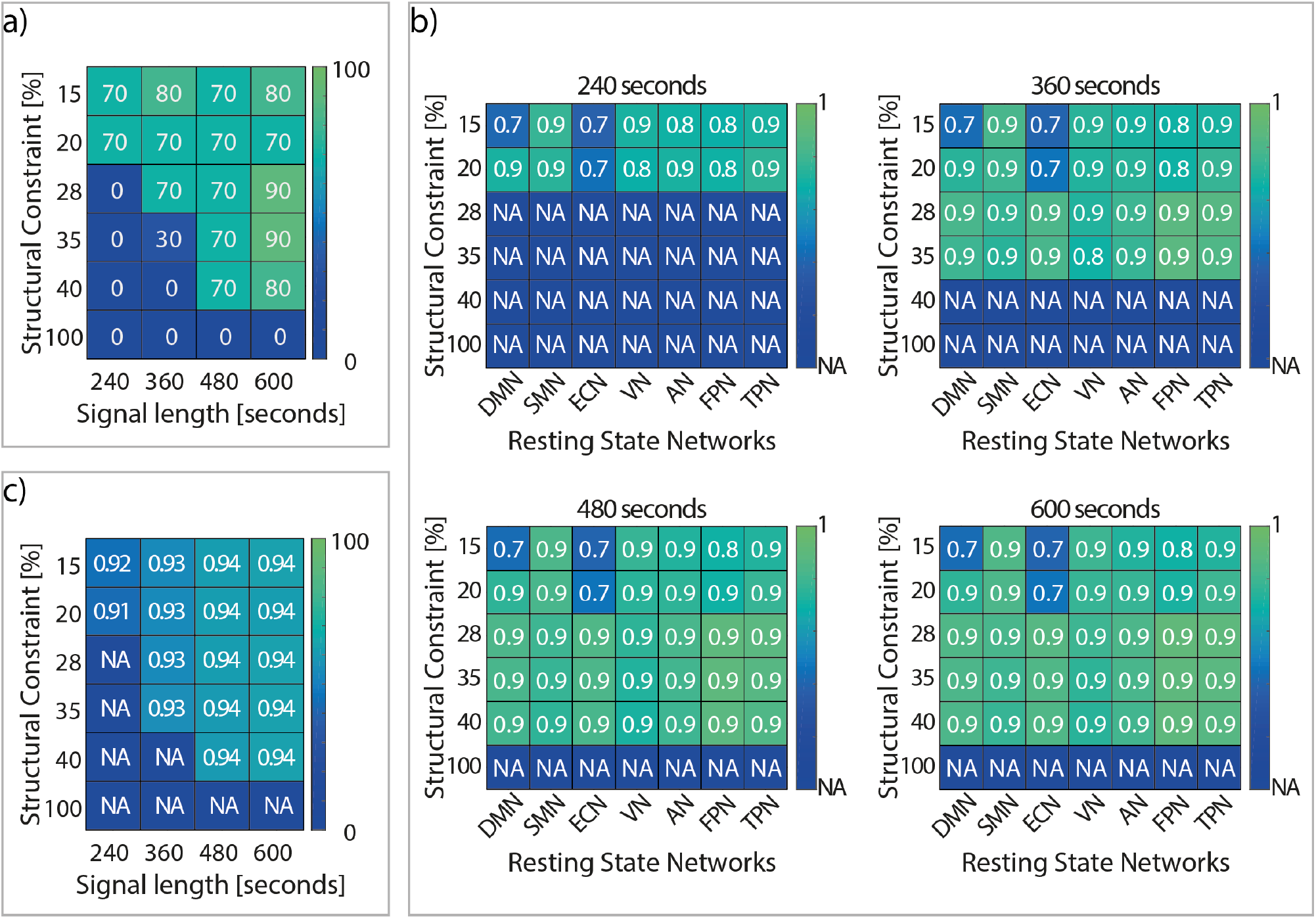
a) Percentage of success in detecting number of RSNs.b) Average MSS of RSNs, varying the percentage of connections and the signal’s length. c) MSS of the estimated transition matrices versus the design transition matrix.

HsMM-MAR estimated explicitly the duration of the states through a probability distribution. The states were designed with a lognormal duration distribution. We calculated the cumulative distribution function (*CDF*), of the state durations, to evaluate the accuracy for estimating the state duration’s probability distribution. Figure 9a) plots the function 1–*CDF*, estimated for the 600 second signals, varying the percentage of structural constraint for each RSN. The x-axis was in logarithmic scale. The black dashed lines indicated the cumulative distribution function obtained with the performance test (analysis performed with G-Test, refer section xx), defining the reliability range. Distributions, outside the reliability range, were considered different from the duration distribution of the simulated signals. The duration probability, of each network, did not present significant differences with the design. Figure 9c) shows the KL divergence between the recovered and design duration distributions for the RSNs, with the 600 seconds signals and varying the structural constraint. The black dashed line indicated the critical KL distance value. All the divergences were under the critical threshold. We assessed the computation time needed for estimations. The HsMM-MAR was forced for detecting seven states to guarantee equality of conditions in each iteration. Figure 9b) illustrates the estimation computation time for the different structural constraints, with respect to the reference structural constraint (28%), for different signal lengths. The computation time increased with structural constraints allowing a higher percentage of anatomical connections.

**Figure 9.**
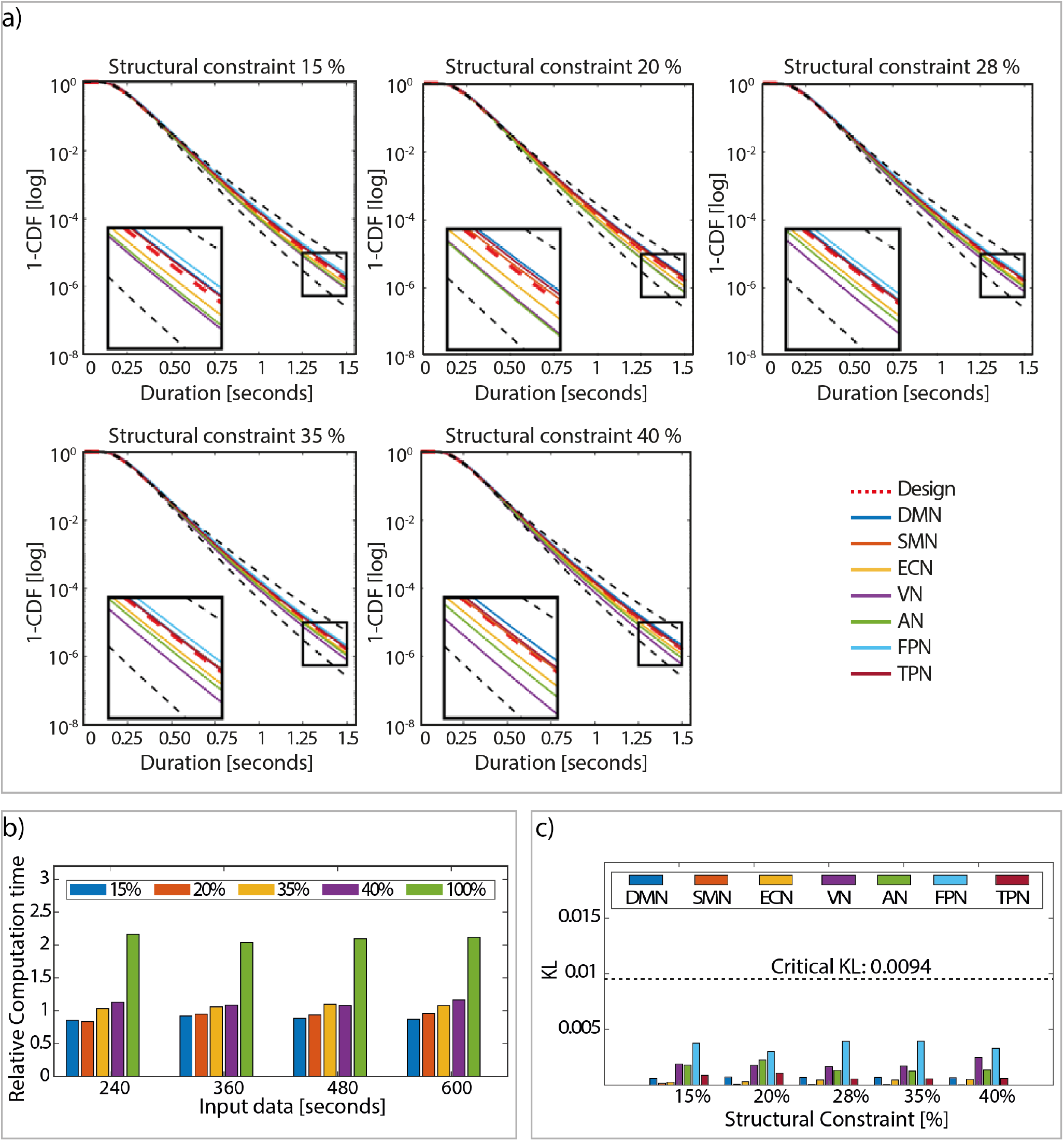
a) The plots display 1 – CDF for states of the the seven RSNs, where CDF is the cumulative distribution function of the state duration, estimated by the algorithm, varying the structural constraint for a signal size of 600 seconds. The red dashed lines indicate the design CDF, the continuous lines show the CDF for the different RSNs, and the the black dashed lines delimit the reliability area. The y-axis is on a logarithmic scale. b) Estimation computation time for the different structural constraints with respect to the estimation time, using a 28% structural constraint. c) KL divergence between the design duration distribution of the states for each RSN and the duration distributions calculated from the parameters estimated by the algorithm.

**Figure 10.**
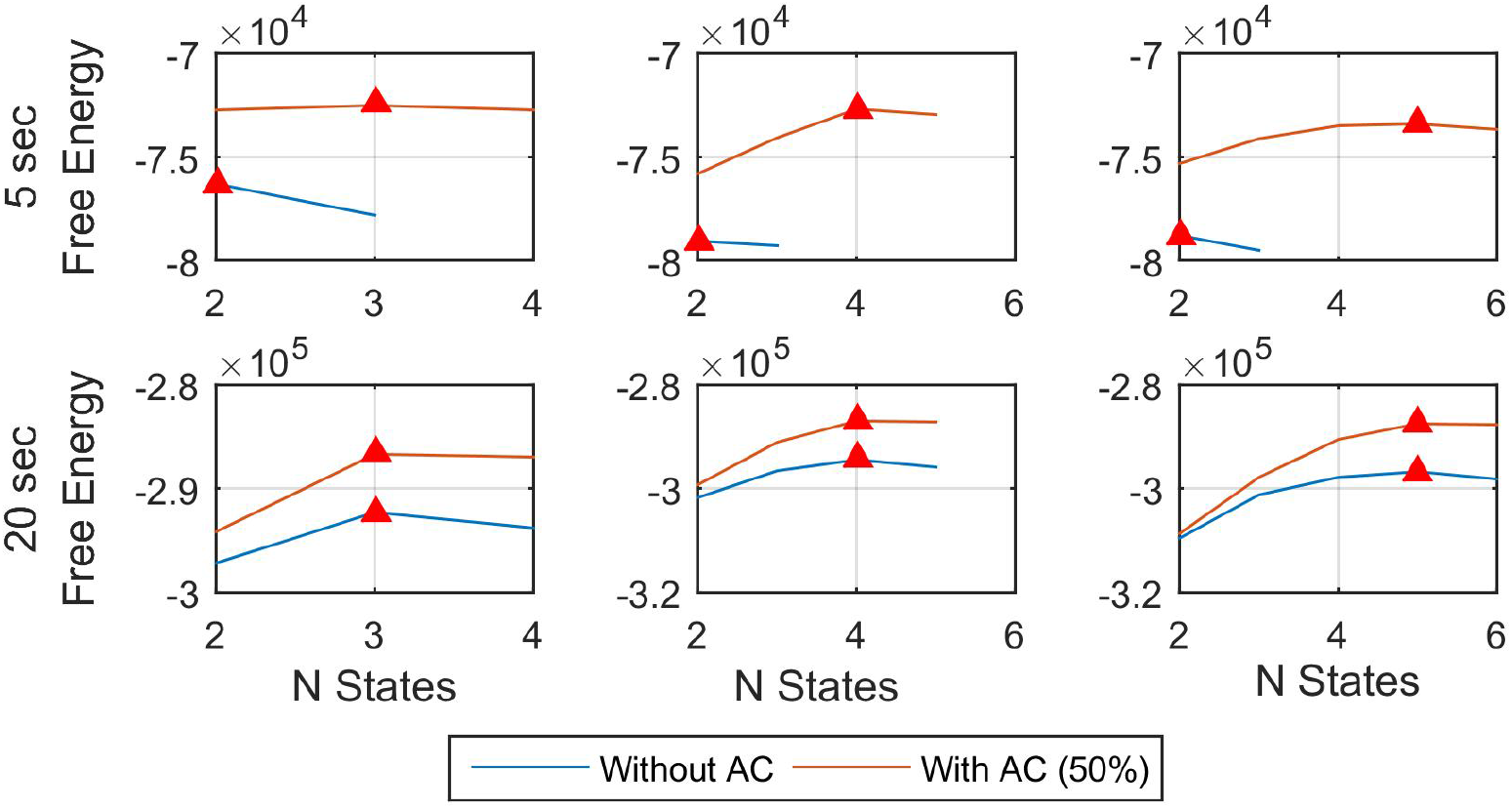
Free Energy convergence curve in training: The estimated number of states corresponds to the maximum of the Free Energy function, in this case marked with the figure of a red triangle. Each graph shows the curve with the model with and without the use of RE.

## 4. DISCUSSION AND CONCLUSION

We presented an efficient method for the analysis of the dynamics of the functional brain signals. The temporal dynamics of brain networks are modelled with a generative HsMM model that allows the modelling of state duration through probability distributions. In addition, each network is characterized by a MAR, enabling the spectral analysis of the network and the temporal interactions between its ROIs. The MARs were constrained using anatomical connectivity and the connection delay between ROIs. These constraints were extracted using diffusion MRI data from the HCP database. Both, anatomical connectivity and the connection delays were represented together using a sparse matrix containing the connection delay between ROIs. The performance of the algorithm was first evaluated using a simple simulation with 10 nodes to validate the correct implementation of the model. Next, we used a more complex simulation based on 62 anatomical ROIs and the 7 most reported RSNs. In both experiments, we assessed the performance of the algorithm for detecting the number of networks, the state duration distributions and the sequence of states for the simulated signals, using different structural constraints and signal length.

The first experiment validated the correct functioning of the model implementation, able to integrate connectivity constraint and connection delay information to the HsMM-MAR model. It is verified that the model is capable of recovering the number of correct states using free energy curve and recover the other parameters correctly.

The second experiment demonstrated that the structural constraint with low percentage of connections enabled a faster convergence of the algorithm even for short signals. On the other hand, when we used all the connections (100% of connections), the algorithm was not able to estimate the correct numbers of RSNs, for the simulated signal lengths. Note that a 100% of structural constraint contains all the connections but only one delay per connection, which is still a sparse matrix. If we do not use structural constraint, the delay matrix is full, and the algorithm is not computationally feasible. Compared to the 100% of structural constraint, the reference connectivity (28% of connections) reduces the complexity of the model, requiring shorter signals. In addition, it reduces the computational cost by around 2 times.

Furthermore, the state duration parameters were recovered with high accuracy when the algorithm correctly detected the number of states. For these cases, the estimated transition probability matrices had a MSS with respect to the design matrices greater than 0.91. In addition, the KL divergence and the G-test between the recovered and design duration distributions, showed that the state duration parameters were obtained with a wide confidence interval for all RSNs.

The second experiment also demonstrated the robustness of the algorithm, since over a minimum signal length for a given constraint, it showed a stable behavior.

Functional brain data are time series, therefore it can be modelled by a Multivariate Autoregressive Regression (MAR) model. There are other data driven methods that can be used to analyze time series. In (Bakhshayesh et al., 2019) compares different data-driven measures methods on simulated data by using three systems (Hénon maps, MARs and simulated EEG). They reported that no measure has superior performance. However, MARs are useful because they perform a spectral analysis of the signal. Several studies have used these models to analyze fMRI signals. (Penny and Roberts et al. 2002) were the first group to apply MAR for the modelling of fMRI signals. A multi-scale generative model for EEG was proposed in (Friston, 2007), which assumes that the mesostates can be modeled by a MAR model. However, they characterize the mesostates as stationary. Therefore (Olier, 2013) proposed the addition of a level of flexibility by using temporal clusters characterized by MAR models. Nevertheless, with this approach it is not possible to obtain a transition probability between mesostates. Subsequently, (Fukushima, 2015) proposed a method for source reconstruction with a MAR model representing interactions among brain regions, defining the model order based on the connection lag.

Previous studies dis not consider the dynamics of brain activity in the estimation of the models. Using a Hidden Markov Model (HMM) it is possible to define the observation model as a MAR, as proposed by (Vidaurre et al., 2016). However, in HMM the distributions of state durations are geometric, assigning high probabilities to faster changes among states. In this regard, (Wegner et al., 2017) demonstrated that the Auto Information Function (AIF) of a Markov process differs critically from an AIF of the real data. Hence, Hidden semi-Markov Model (HsMM) arises as an excellent tool to model the distribution of state duration ((Trujillo-Barreto et al., 2019)). However, MAR observation model may be excessively complex and computationally unapproachable.

The number of parameters to estimate can be calculated as (n*n*d), where n is the number of nodes and d the order of the MAR. In addition, complex models can suffer from overfitting. This occurs when the model cannot correctly generalize from the training data and performs poorly for the testing data (Ying, 2019). In some cases, overfitting occurs by building a model with little data for training with respect to the parameters to be estimated (Bishop, 2006; Ying, 2019). In extreme cases, the model may be unidentifiable, when the training set is not large enough to converge to a single solution. In these cases, it is possible to apply model constraints to reduce the parameters to be estimated (Godfrey and DiStefano, 1985). For this reason, we proposed an HsMM-MAR-AC approach that allows a considerable reduction in the parameters to be estimated.

Functional integration between different areas in the brain are mediated by white matter (WM) connections (Sotiropoulos and Zalesky, 2019). Koch et al. (2002) found that high structural connectivity involves high functional connectivity. Later, several studies have reported that functional dynamics are a reflection of structural connectivity (Damoiseaux and Greicius, 2009). These findings suggest that anatomical connectivity can be used as a constraint in the study of functional connectivity. In works such as Penny et al. (2005), Fukushima et al. (2013) and Fukushima et al. (2015) structural connectivity is applied as a constraint to MAR models of high complexity. Recently, in (Crimi et al, 2021), an autoregressive model was used for studying the association between structural and functional brain activity, by using a multi-lag autoregressive constrained for the structural constraint. They studied the relationship between structure and functionality of brain. They found that the use of effective connectivity better describes the functional interactions, maintaining the original structural organization and discriminating between cases and controls.

The authors used the Akaike information criterion (Akaike et al, 1974) for defining the MAR order. On the other hand, Fukushima et al. (2015) applied an arbitrary threshold to define the structural constraint. In our work, we use the average weighted connectivity matrix (equation9) for thresholding the anatomical constraint (AC) and the maximum fiber length.

The anatomical constraint discards the weakest connections, which may cause loss of some RSN connections. In this work we used a reference threshold, where all the RSN connections are preserved, leading to a connectivity matrix with 28% of connections. Most restrictive structural constraints eliminate some long (a higher proportion of long connections than short) connections. This seems to suggest that the connection weights are stronger for short connections. However, the correlation between the delay matrix and the average weighted connectivity matrix has a low correlation (−0.30). This occurs because there are very long connections that are not important in RSNs, which have low connection weights. In fact, the 15% and 20% connection constraints eliminate some short connections of the DMN and ECN, while maintaining the long connections of all RSNs.

When removing connections of some RSNs, the MSS between the simulated and estimated networks is lower for the affected networks. However, even if connections are lost, it is possible to obtain with high accuracy both the temporal dynamics of the networks and the probability of state transition. In addition, shorter signals are needed to obtain the dynamics, what is relevant since functional activity is generally measured by short signals. In many cases, it is necessary to concatenate signals from several subjects. We may avoid loss of connections by using a 100% structural constraint, however the accuracy for recovering the numbers of RSNs decreases to 40%. Besides, the MSS between the simulated and estimated networks decreases, requiring larger signals to obtain the same performance as for constraints with fewer connections. The best performance is obtained for the reference connectivity (28%), which demonstrates the relevance of validating a method to assess the accuracy for obtaining the optimal structural constraint. It is possible to use only the RSN connections. In this scenario the performance of the algorithm should be better. However, to avoid introducing a bias we were as objective as possible to define the thresholds of the structural constraints guided by the biological information (connection weight).

In real data, it is possible that the delay of connections is variable among different people due to diseases (Berman, 2020), aging, or the natural development of brain connections, resulting from their experiences. However, (Berman, 2019) modeled the conduction delays in the corpus callosum and did not find differences with age. In our work, we used a single delay for each connection, which may be a simplistic approach. Nevertheless, we used 5 ms of temporal resolution, thus we may assume that the delay variability is absorbed by the sample rate.

In the connection delay representation, a significant aspect to consider is the number of ROIs to characterize the networks. If more ROIs are used, we may obtain better descriptions of the interactions, however, it may be necessary to use very restrictive structural constraints due to increase of the model complexity. In any case, we used a ROI atlas extensively used in the literature. Future work could explore different atlases and evaluate the impact of the atlas selection.

The results prove the impact of the AC on the performance of the algorithm. The accuracy of the estimation of the number of states decreases with more relaxed AC. To obtain a better performance would require longer signals and estimation times. In any case, AC turns the complexity of the algorithm computationally approachably. For more restrictive constraints, that lead to the loss of some RSN connections, the algorithm is still able to recognize all 7 RSNs, what shows the robustness of the algorithm to errors in the definition of structural connectivity. Furthermore, the accuracy in the estimation of the state duration distributions demonstrate a correct estimation of the signal dynamics. The proposed method enables an efficient and feasible tool to analyze brain dynamics.

## AUTHOR CONTRIBUTIONS

**Hernan Hernandez Larzabal:** Conceptualization, Methodology, Software, Investigation, Writing – original draft, Writing – review & editing, Visualization. **David Araya:** Conceptualization, Methodology, Software, Investigation, Writing – original draft, Writing – review & editing, Visualization. **Liset Gonz** á **lez Rodr** í **guez:** Methodology, Investigation, Writing – original draft, Writing – review & editing **Claudio Rom**á**n:** Resources, Data pre-processing **Nelson Trujillo-Barreto:** Conceptualization, Methodology, Formal analysis, Writing - original draft, Writing - review & editing, Supervision **Pamela Guevara:** Conceptualization, Methodology, Formal analysis, Writing - original draft, Writing - review & editing, Supervision **Wael El-Deredy:** Conceptualization, Methodology, Formal analysis, Writing - original draft, Writing - review & editing, Supervision, Project administration.

## DATA AVAILABILITY STATEMENT

All data used in this work is from the Human Connec-tome Project (HCP) www.humanconnectome.org, which is publicly available to researchers who agree to the data use terms www.humanconnectome.org/study/hcp-young-adult/data-use-terms. All HCP data may be downloaded through the ConnectomeDB db.humanconnectome.org.

## ACKNOWLEDGMENTS

HCP Data were provided by the Human Connectome Project, WUMinn Consortium (Principal Investigators: David Van Essen and Kamil Ugurbil; 1U54MH091657) funded by the 16 NIH Institutes and Centers that support the NIH Blueprint for Neuroscience Research; and by the McDonnell Center for Systems Neuroscience at Washington University.

