## Supplementary for "Efficient estimation of time-dependent functional connectivity using Structural Connectivity constraints"

### SUPPLEMENTARY MATERIAL

#### Emission Model Upgrade Rules

1. W Coefficient MAR

$$log(q(w_{nl_n}^{(k)})) = log(N(w_{nl_n}^{(k)}|\bar{w}_{nl_n}^{(k)}, {\Upsilon_n^{(k)}}^{-1})) \tag{1}$$

With:

$$\bar{w}_{nl_n}^{(k)} = \sum_{t=1}^{T} \gamma_t^{(k)} * \bar{\phi}_n^{(k)} * {\Upsilon_n^{(k)}}^{-1} * {x_{l_n t}}^T * y_{nt} \tag{2}$$

$$\Upsilon_n^{(k)} = (\sum_{t=1}^{T} \gamma_t^{(k)} * \bar{\lambda}_{nl_n}^{(k)} * \bar{\phi}_n^{(k)} * x_{l_n t} * {x_{l_n t}}^T)^{-1} \tag{3}$$

Where $\bar{w}_{nl_n}^{(k)}$ and $\Upsilon_n^{(k)}$ correspond to the mean and Precision of Normal distribution.

2. MAR coefficient Precision

$$log(q(\lambda_{nj}^{(k)})) = Gam(\lambda_{nj}^{(k)}|b_{nj}^{(k)}, c_{nj}^{(k)}) \tag{4}$$

With:

$$b_{nj}^{(k)} = \hat{b}_{nj} + \frac{1}{2} \tag{5}$$

$$c_{nj}^{(k)} = (\frac{1}{\hat{c}_{nj}} + \frac{1}{2} * ({\bar{w}_n^{(k)}}^T * \bar{w}_n^{(k)} + Tr(\Upsilon_n^{(k)})))^{-1} \tag{6}$$

Where $b_{nj}^{(k)}$ and $c_{nj}^{(k)}$ correspond to the shape and scale of Gamma distribution.

3. Observation Noise Precision

$$log(q(\phi_n^{(k)})) = Gam(\phi_n^{(k)}|e_n^{(k)}, f_n^{(k)}) \tag{7}$$

With:

$$e_n^{(k)} = \hat{e}_n + \frac{\sum_{t=1}^{N} \gamma_t^{(k)}}{2}$$

$$f_n^{(k)} = \{\frac{1}{\hat{f}_n} + \sum_{t=1}^{N} \frac{1}{2} * \gamma_t^{(k)} * ((y_{nt} - {\bar{w}_{nl_n}^{(k)}}^T * x_{l_n t})^2 + {x_{l_n t}}^T * {\Upsilon_n^{(k)}}^{-1} * x_{l_n t})\}^{-1} \quad (8)$$

Where $e_n^{(k)}$ and $f_n^{(k)}$ correspond to the shape and scale of Gamma distribution.

### ADITIONAL FIGURES

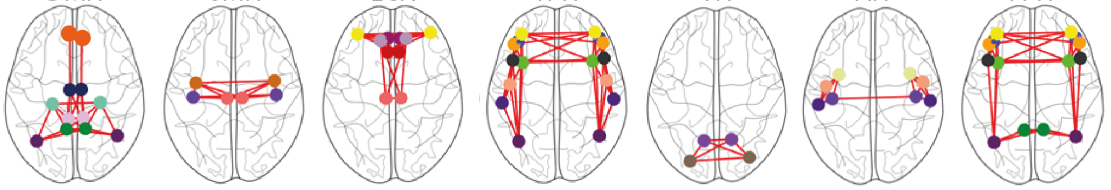

| | DMN | SMN | ECN | TPN | VN | AN | FPN |
|---|---|---|---|---|---|---|---|
| CAM | | | X | | | | |
| CMF | | | | X | | | X |
| InfP | X | | | X | | | X |
| InfT | | | | | | | |
| IC | X | | | | | | |
| LOcc | | | | | X | | |
| LinG | | | | | X | | |
| MOrb | X | | | | | | |
| MT | | | | | | X | |
| PH | X | | | | | | |
| PCen | | X | X | | | | |
| POper | | | | X | | | X |
| POrb | | | | X | | | X |
| PTria | | | | X | | | X |
| PostC | | X | | | | X | |
| PostCin | X | | | | | | |
| Prec | | X | | | | | |
| PCun | X | | | X | | | X |
| RAC | | | X | | | | |
| RMF | | | X | X | | | X |
| SupF | | | X | | | | |
| SupT | | | | | | X | |
| Ins | | | | | | X | |

**Figure 1.** **[this figure needs color in printing]** Nodes of the Resting State Networks [RSN] based on the Desikan-Killiany atlas thresholded at 28% of all anatomical connections as described in **??**, which conserved all RSNs. Abreviations: CAM (Caudal Anterior Cingulate), CMF (Caudal Middle Frontal), CN (Cuneus), ENT (Entorhinal), FUS (Fusiform), InfP (Inferior Parietal), InfT (Inferior Temporal), IC (Isthmus Cingulate), LOcc (Lateral Occipital), LOrb (Lateral Orbitofrontal), LinG (Lingual), MOrb (Medial Orbitofrontal), MT (Middle Temporal), PH (Parahippocampal), PCen (Paracentral), POper (Pars Opercularis), POrb (Pars Orbitalis), PTria (Pars Triangularis), Pcal (Pericalcarine), PostC (Postcentral), PostCin (Posterior Cingulate), Prec (Precentral), PCun (Precuneus), RAC (Rostral Anterior Cingulate), RMF (Rostral Middle Frontal), SupF (Superior Frontal), SupP (Superior Parietal), SupT (Superior Temporal), Supra (Supramarginal), TT (Transverse Temporal), Ins (Insula).

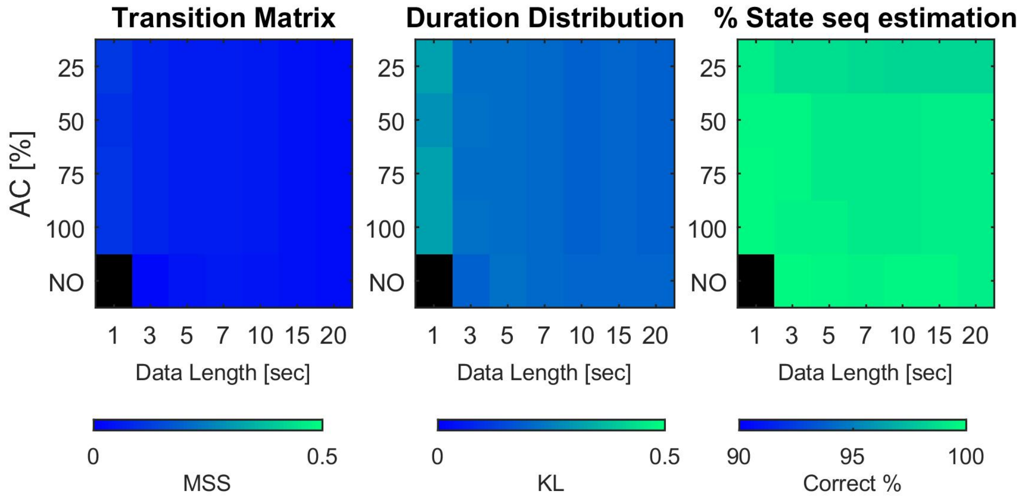


**Figure 2.** [**this figure needs color in printing**] In the three graphs, the Y axis indicates the % of AC used and the X axis the duration in second. Figure A shows the recovery performance of the transition matrix by means of the MSS distance between the real matrix and estimated one. The range of this metric is from 0 to 1 and it was obtained only for those cases where the number of states was correctly determined. The black color indicates that it is not possible to calculate this metric because there are no cases where it converges to the correct number of states. In most cases the recovery is very good, resulting in values around 0.Figure B shows the recovery performance of the durations distribution by meansof the KL distance between the real distribution and the estimated one. In most cases the recovery is very good, resulting in values around 0.1.Figure C shows the % recovery of the states sequence. In most cases the recovery is very good, resulting in values around 98%.
